# Simulating a Primary Visual Cortex at the Front of CNNs Improves Robustness to Image Perturbations

**DOI:** 10.1101/2020.06.16.154542

**Authors:** Joel Dapello, Tiago Marques, Martin Schrimpf, Franziska Geiger, David D. Cox, James J. DiCarlo

## Abstract

Current state-of-the-art object recognition models are largely based on convolutional neural network (CNN) architectures, which are loosely inspired by the primate visual system. However, these CNNs can be fooled by imperceptibly small, explicitly crafted perturbations, and struggle to recognize objects in corrupted images that are easily recognized by humans. Here, by making comparisons with primate neural data, we first observed that CNN models with a neural hidden layer that better matches primate primary visual cortex (V1) are also more robust to adversarial attacks. Inspired by this observation, we developed VOneNets, a new class of hybrid CNN vision models. Each VOneNet contains a fixed weight neural network front-end that simulates primate V1, called the VOneBlock, followed by a neural network back-end adapted from current CNN vision models. The VOneBlock is based on a classical neuroscientific model of V1: the linear-nonlinear-Poisson model, consisting of a biologically-constrained Gabor filter bank, simple and complex cell nonlinearities, and a V1 neuronal stochasticity generator. After training, VOneNets retain high ImageNet performance, but each is substantially more robust, outperforming the base CNNs and state-of-the-art methods by 18% and 3%, respectively, on a conglomerate benchmark of perturbations comprised of white box adversarial attacks and common image corruptions. Finally, we show that all components of the VOneBlock work in synergy to improve robustness. While current CNN architectures are arguably brain-inspired, the results presented here demonstrate that more precisely mimicking just one stage of the primate visual system leads to new gains in ImageNet-level computer vision applications.

## 1 Introduction

For the past eight years, convolutional neural networks (CNNs) of various kinds have dominated object recognition [1, 2, 3, 4], even surpassing human performance in some benchmarks [5]. However, scratching beneath the surface reveals a different picture. These CNNs are easily fooled by imperceptibly small perturbations explicitly crafted to induce mistakes, usually referred to as adversarial attacks [6, 7, 8, 9, 10]. Further, they exhibit a surprising failure to recognize objects in images corrupted with different noise patterns that humans have no trouble with [11, 12, 13]. This remarkable fragility to image perturbations has received much attention in the machine learning community, often from the perspective of safety in real-world deployment of computer vision systems [14, 15, 16, 17, 18, 19, 20, 21, 22]. As these perturbations generally have no perceptual alignment with the object class [23], the failures suggest that current CNNs obtained through task-optimization end up relying on visual features that are not all the same as those used by humans [24, 25]. Despite these limitations, some CNNs have achieved unparalleled success in partially explaining neural responses at multiple stages of the primate ventral stream, the set of cortical regions underlying primate visual object recognition [26, 27, 28, 29, 30, 31, 32].

### How can we develop CNNs that robustly generalize like human vision?

Incorporating biological constraints into CNNs to make them behave more in line with primate vision is an active field of research [33, 34, 35, 36, 37, 38, 39, 40, 41, 42, 43, 32]. Still, no neurobiological prior has been shown to considerably improve CNN robustness to both adversarial attacks and image corruptions in challenging real-world tasks such as ImageNet [44]. Here, we build on this line of work, starting with the observation that the ability of each CNN to explain neural response patterns in primate primary visual cortex (V1) is strongly correlated with its robustness to imperceptibly small adversarial attacks. That is, the more biological a CNN’s "V1" is, the more adversarially robust it is.

Inspired by this, we developed *VOneNets*, a new class of hybrid CNNs, containing a biologically-constrained neural network that simulates primate V1 as the front-end, followed by an off-the-shelf CNN back-end trained using standard methods. The V1 front-end, *VOneBlock*, is based on the classical neuroscientific linear-nonlinear-Poisson (LNP) model, consisting of a fixed-weight Gabor filter bank (GFB), simple and complex cell nonlinearities, and neuronal stochasticity. The VOneBlock outperforms all standard ImageNet trained CNNs we tested at explaining V1 responses to naturalistic textures and noise samples. After training, VOneNets retain high ImageNet performance, but are substantially more robust than their corresponding base models, and compete with state-of-the-art defense methods on a conglomerate benchmark covering a variety of adversarial images and common image corruptions. Importantly, these benefits transfer across different architectures including ResNet50 [4], AlexNet [1], and CORnet-S [32]. We dissect the VOneBlock, showing that all properties work in synergy to improve robustness and that specific aspects of VOneBlock circuitry offer robustness to different perturbation types. Notably, we find that neuronal stochasticity plays a large role in the white box adversarial robustness, but that stochasticity alone is insufficient to explain our results—neuronal stochasticity interacts supralinearly with the VOneBlock features to drive adversarial robustness. Finally, as a large percentage of this robustness remains even when we remove stochasticity only during the adversarial attack, we conclude that training with stochasticity at the VOneBlock level leads the downstream layers to learn more robust representations.

Model weights and code are available at https://github.com/dicarlolab/vonenet.

### 1.1 Related Work

#### V1 modeling

Since the functional characterization of simple and complex cell responses by Hubel and Wiesel [45], modeling V1 responses has been an area of intense research. Early approaches consisting of hand-designed Gabor filters were successful in predicting V1 simple cell [46] and complex cell [47] responses to relatively simple stimuli. These models were later improved with the addition of further nonlinear operations, such as normalization and pooling, to account for extra-classical functional properties [48, 49]. Early hierarchical models of object recognition, which incorporated these type of V1 models as their early layers, were used with some success in modeling neuronal responses in the ventral stream and object recognition behavior [50, 51]. Generalized LNP models expanded the V1 model class by allowing a set of fitted excitatory and suppressive spatial filters to be nonlinearly combined [52], and subunit models introduced two sequential linear-nonlinear (LN) stages for fitting V1 responses [53]. Recently, both task-optimized and data-fitted CNNs were shown to narrowly beat a GFB model in predicting V1 responses, further validating that multiple LN stages may be needed to capture the complexity of V1 computations [30].

#### Model robustness

Much work has been devoted to increasing model robustness to adversarial attacks [54, 17, 15, 55, 56, 40, 57], and to a lesser extent common image corruptions [13, 58, 59]. In the case of adversarial perturbations, the current state-of-the-art is adversarial training, where a network is explicitly trained to correctly classify adversarially perturbed images [17]. Adversarial training is computationally expensive [60, 61], known to impact clean performance [62], and overfits to the attack constraints it is trained on [63, 64]. Other defenses involve adding noise either during training [59], inference [65, 19], or both [15, 66, 67]. In the case of stochasticity during inference, Athalye et. al. demonstrated that fixing broken gradients or taking the expectation over randomness often dramatically reduces the effectiveness of the defense [68]. In a promising demonstration that biological constraints can increase CNN robustness, Li et. al. showed that biasing a neural network’s representations towards those of the mouse V1 increases the robustness of grey-scale CIFAR [69] trained neural networks to both noise and white box adversarial attacks [40].

## 2 Adversarial Robustness Correlates with V1 Explained Variance

The susceptibility of current CNNs to be fooled by imperceptibly small adversarial perturbations suggests that these CNNs rely on some visual features not used by the primate visual system. Are models that better explain neural responses in the macaque V1 more robust to adversarial attacks? We analyzed an array of publicly available neural networks with standard ImageNet training [70] including AlexNet [1], VGG [3], ResNet [4], ResNeXt [71], DenseNet [72], SqueezeNet [73], ShuffleNet [74], and MnasNet [75], as well as several ResNet50 models with specialized training routines, such as adversarial training with *L*_∞_ (‖*δ*‖_∞_ = 4/255 and ‖*δ*‖_∞_ = 8/255) and *L*_2_ (‖*δ*‖_2_ = 3) constraints [76], and adversarial noise combined with Stylized ImageNet training [59].

For each model, we evaluated how well it explained the responses of single V1 neurons evoked by given images using a standard neural predictivity methodology based on partial least square regression (PLS) [31, 32]. We used a neural dataset with 102 neurons and 450 different 4deg images, consisting of naturalistic textures and noise samples [77]. Explained variance was measured using a 10-fold cross-validation strategy. Similarly to other studies, we considered the field-of-view of all models to span 8deg [31, 30] (see Supplementary Section A for more details). To evaluate the adversarial robustness of each model in the pool, we used untargeted projected gradient descent (PGD) [17], an iterative, gradient-based white box adversarial attack with *L*_∞_, *L*_2_, and *L*_1_ norm constraints of ‖*δ*‖_∞_ = 1/1020, ‖*δ*‖_2_ = 0.15, and ‖*δ*‖_1_ = 40, respectively. For each norm, the attack strength was calibrated to drive variance in performance amongst non-adversarially trained models, resulting in perturbations well below the level of perceptibility (see Supplementary Section B.1 for more details).

We found that accuracy under these white box adversarial attacks has a strong positive correlation with V1 explained variance (Fig. 1, r=0.85, p=2.1E-9, n=30). Notably, adversarially trained ResNet50 models [76], which were hardly affected by these attacks, explained more variance in V1 neural activations than any non-adversarially trained neural network. The correlation was not driven by the CNNs’ clean ImageNet performance since it was even more pronounced when the white box accuracy was normalized by the clean accuracy (r=0.94, p=7.4E-15, n=30) and was also present when removing the adversarially trained models (r=0.73, p=1.78E-5, n=27). While increasing the perturbation strength rapidly decreases the accuracy of non-adversarially trained models, the described correlation was present for a wide range of attack strengths: when the perturbation was multiplied by a factor of 4, greatly reducing the variance of non-adversarially trained models, white box accuracy was still significantly correlated with explained variance in V1 (r=0.82, p=1.17E-8, n=30).

**Figure 1:**
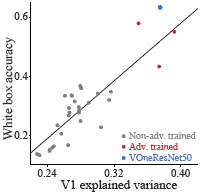
CNNs’ robustness to white box attacks correlates with explained response variance in primate V1. Comparison of top-1 accuracy under white box attacks of low perturbation strengths (average of 3 PGD constraints: ‖*δ*‖_∞_ = 1/1020, ‖*δ*‖_2_ = 0.15, and ‖*δ*‖_1_ = 40) against fraction of explained variance of V1 responses (using PLS regression) for a pool of CNN models. Perturbation strength was chosen to drive variance across model performance. White box accuracy and V1 explained variance are significantly correlated (r=0.85, p=2.1E-9, n=30 CNNs, linear fit shown in gray line). Gray circles, non-adversarially trained trained CNNs (n=27); red circles, adversarially trained ResNet50 models (n=3); blue circle, VOneResNet50 (not included in correlation).

## 3 VOneNet: a Hybrid CNN with a V1 Neural Network Front-End

Inspired by the strong correlation between V1 explained variance and robustness to white box attacks, we developed the VOneNet architecture. The major characteristic that sets the VOneNet architecture apart is its V1 front-end, the VOneBlock. While most of its layers have parameters learned during ImageNet training, VOneNet’s first block is a fixed-weight, mathematically parameterized CNN model that approximates primate neural processing of images up to and including area V1. Importantly, the VOneBlock components are mapped to specific neuronal populations in V1, which can be independently manipulated or switched off completely to evaluate their functional role. Finally, the VOneBlock can be easily adapted to different CNN base architectures as described below. Here we build VOneNets from three base architectures: ResNet50 [4], CORnet-S [32], and AlexNet [1].

A VOneNet consists of the VOneBlock and a back-end network adapted from a base CNN (Fig. 2). When building a VOneNet from a particular CNN, we replace its first block (typically one stack of convolutional, normalization, nonlinear, and pooling layers, Supplementary Table C.3) by the VOneBlock and a trained transition layer. The VOneBlock matches the replaced block’s spatial map dimensions (56 × 56 for the base CNNs considered) but can have more channels (for the standard model we chose *C*_*V*1_ = 512; see Supplementary Fig. C.2 for an analysis of how the number of channels affects the results). It is followed by the transition layer, a 1 × 1 convolution, that acts as a bottleneck to compress the higher channel number to the original block’s depth. The VOneBlock is inspired by the LNP model of V1 [52], consisting of three consecutive processing stages—convolution, nonlinearity, and stochasticity generator—with two distinct neuronal types—simple and complex cells—each with a certain number of units per spatial location (standard model with *SC*_*V*1_ = *CC*_*V*1_ = 256). The following paragraphs describe the main components of the VOneBlock (see Supplementary Section C for a more detailed description).

**Figure 2:**
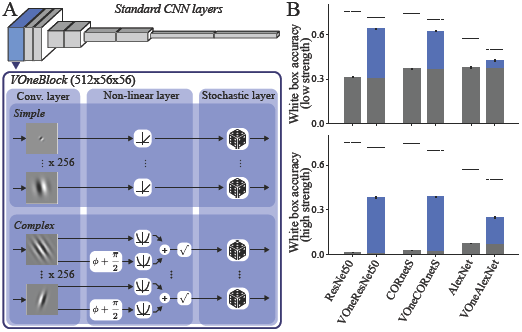
Simulating V1 at the front of CNNs improves robustness to white box attacks. **A** A VOneNet consists of a model of primate V1, the VOneBlock, followed by a standard CNN architecture. The VOneBlock contains a convolutional layer (a GFB with fixed weights constrained by empirical data), a nonlinear layer (simple or complex cell nonlinearities), and a stochastic layer (V1 stochasticity generator with variance equal to mean). **B** Top, comparison of accuracy (top-1) on low strength white box attacks (PGD with ‖*δ*‖_∞_ = 1/1020, ‖*δ*‖_2_ = 0.15, ‖*δ*‖_1_ = 40) between three base models (ResNet50, CORnet-S, and AlexNet) and their correspoding VOneNets. Gray bars show the performance of base models. Blue bars show the improvements from the VOneBlock. Dashed lines indicate the performance on clean images. Bottom, same but for white box attacks of higher strength (PGD with ‖*δ*‖_∞_ = 1/255, ‖*δ*‖_2_ = 0.6, ‖*δ*‖_1_ = 160). VOneNets show consistently higher white box accuracy even for perturbation strengths that reduce the performance of the base models to nearly chance. Error-bars represent SD (n=3 seeds).

### Biologically-constrained Gabor filter bank

The first layer of the VOneBlock is a mathematically parameterized GFB with parameters tuned to approximate empirical primate V1 neural response data. It convolves the RGB input images with Gabor filters of multiple orientations, sizes, shapes, and spatial frequencies, in a 56 × 56 spatial map. To instantiate a VOneBlock, we randomly sample *C*_*V*1_ values for the GFB parameters according to empirically observed distributions of preferred orientation, peak spatial frequency, and size/shape of receptive fields [78, 79, 80]. The VOneBlock keeps color processing separate, with each Gabor filter convolving a single color channel from the input image. The resulting set of spatial filters is considerably more heterogeneous than those found in the first layer of standard CNNs, and better approximates the diversity of primate V1 receptive fields (Supplementary Fig. C.1).

### Simple and complex cells

While simple cells were once thought to be an intermediate step for computing complex cell responses, it is now known that they form the majority of downstream projections to V2 [81]. For this reason, the VOneBlock nonlinear layer has two different nonlinearities that are applied to each channel depending on its cell type: a rectified linear transformation for simple cells, and the spectral power of a quadrature phase-pair for complex cells.

### V1 stochasticity

A defining property of neuronal responses is their stochasticity—repeated measurements of a neuron in response to nominally identical visual inputs results in different spike trains. In awake monkeys, the mean spike count (averaged over many repetitions) depends on the presented image, and the spike train for each trial is approximately Poisson: the spike count variance is equal to the mean [82]. To approximate this property of neuronal responses, we add independent Gaussian noise to each unit of the VOneBlock, with variance equal to its activation. Before doing this, we apply an affine transformation to the units’ activations so that both the mean stimulus response and the mean baseline activity are the same as those of a population of primate V1 neurons measured in a 25ms time-window (mean stimulus response and spontaneous activity of 0.324 and 0.073 spikes, respectively; see Supplementary Table C.2 and Fig. C.2 for other time-windows). Like in the brain, the stochasticity of the VOneBlock is always on, during both training and inference.

### V1 explained variance

The VOneBlock was not developed to compete with state-of-the-art data-fitting models of V1 [53, 30]. Instead we used available empirical distributions to constrain a GFB model, generating a neuronal space that approximates that of primate V1. Despite its simplicity, the VOneBlock outperformed all tested standard ImageNet trained CNNs in explaining responses in the V1 dataset used, and with an explained variance of 0.375±0.006, came right in the middle of the range of the adversarially trained CNNs (Fig. 1, blue circle). On the surface, it seems that these results are at odds with Cadena et al which showed that a task-optimized CNN marginally outperformed a GFB model in explaining V1 responses [30]. However, our GFB has parameters constrained by empirical data resulting in a better model of V1; when we use the parameters of the GFB in Cadena et al, we obtained a much lower explained variance (0.296±0.005). Our results suggest that a well-tuned GFB and adversarially trained CNNs are currently the best models in predicting primate V1 responses.

## 4 Results

### 4.1 Simulating a V1 at the front of CNNs improves robustness to white box attacks

To evaluate the VOneNets’ robustness to whitebox adversarial attacks, we used untargeted PGD with *L*_∞_, *L*_2_, and *L*_1_ norm constraints of ‖*δ*‖_∞_ ∈ [1/1020, 1/255], ‖*δ*‖_2_ ∈ [0.15, 0.6], and ‖*δ*‖_1_ ∈ [40, 160]. All VOneNets were attacked end-to-end, with gradients propagated through the VOneBlock to the pixels. Because the VOneBlock is stochastic, we took extra care when attacking VOneNets. We use the reparameterization trick when sampling VOneBlock stochasticity, allowing us to keep the gradients intact [83]. Further, to combat the effects of noise in the gradients, we adapted our attack such that at every PGD iteration, we take 10 gradient samples and move in the average direction to increase the loss [68] (Supplementary Table B.1). To verify the effectiveness of our attacks, we used controls suggested by Carlini et. al. 2019 [84]; in particular, we show that increasing the magnitude of the norm of the attack monotonically decreases VOneNet accuracy, eventually reaching 0% accuracy (Supplementary Fig. B.1). Additionally, increasing the number of PGD steps from 1 to many strictly increases the effect of the attack under a given norm size (Supplementary Fig. B.2). Both of these indicate that VOneNet gradients contain useful information for the construction of adversarial examples. For more details and controls, see Supplementary Sections B.1 and B.2.

We found that simulating a V1 front-end substantially increased the robustness to white box attacks for all three base architectures that we tested (ResNet50, CORnet-S, and AlexNet) while leaving clean ImageNet performance largely intact (Fig. 2 B). This was particularly evident for the stronger perturbation attacks, which reduced the accuracy of the base models to nearly chance (Figs. 2 B bottom). This suggests that the VOneBlock works as a generic front-end, which can be transferred to a variety of different neural networks as an architectural defense to increase robustness to adversarial attacks.

### 4.2 VOneNets outperform state-of-the-art methods on a composite set of perturbations

We then focused on the ResNet50 architecture and compared VOneResNet50 with two state-of-the-art training-based defense methods: adversarial training with a ‖*δ*‖_∞_= 4/255 constraint (AT_*L*∞_) [76], and adversarial noise with Stylized ImageNet training (ANT^3×3^+SIN) [59]. Because white box adversarial attacks are only part of the issue of model robustness, we considered a larger panel of image perturbations containing a variety of common corruptions. For evaluating model performance on corruptions we used the ImageNet-C dataset [13] which consists of 15 different corruption types, each at 5 levels of severity, divided into 4 categories: noise, blur, weather, and digital (see Supplementary Fig. B.5 for example images and Supplementary Section B.3 for more details).

As expected, each of the defense methods had the highest accuracy under the perturbation type that it was designed for, but did not considerably improve over the base model on the other (Table 1). While the AT_*L*∞_ model suffered substantially on corruptions and clean performance, the ANT^3×3^+SIN model, the current state-of-the-art for common corruptions, had virtually no benefit on white box attacks. On the other hand, VOneResNet50 improved on both perturbation types, outperforming all the models on perturbation mean (average of white box and common corruptions) and overall mean (average of clean, white box, and common corruptions), with a difference to the second best model of 3.3% and 5.3%, respectively. Specifically, VOneResNet50 showed substantial improvements for all the white box attack constraints (Supplementary Fig. B.1), and more moderate improvements for common image corruptions of the categories noise, blur, and digital (Supplementary Table B.2 and Fig. B.6). These results are particularly remarkable since VOneResNet50 was not optimized for robustness and does not benefit from any computationally expensive training procedure like the other defense methods. When compared to the base ResNet50, which has an identical training procedure, VOneResNet50 improves 18% on perturbation mean and 10.7% on overall mean (Table 1).

**Table 1:**
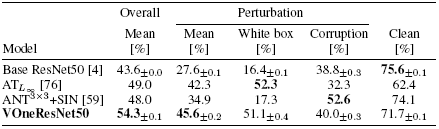
VOneResNet50 outperforms other defenses on perturbation mean and overall mean. Accuracy (top-1) on overall mean, perturbation mean, white box attacks, common corruptions, and clean images for standard ResNet50 and three defense methods: adversarial training, adversarial noise combined with Stylized ImageNet training, and VOneResNet50. Perturbation mean is the average accuracy over white box attacks and common image corruptions. The overall mean is the average accuracy over the two perturbation types and clean ImageNet. Values are reported as mean and SD (n=3 seeds).

### 4.3 All components of the VOneBlock contribute to improved model robustness

Since the VOneBlock was not explicitly designed to increase model robustness, but rather to approximate primate V1, we investigated which of its components are responsible for the increased robustness we observe. We performed a series of experiments wherein six new VOneNet variants were created by removing or modifying a part of the VOneBlock (referred to as VOneBlock variants). After ImageNet training, model robustness was evaluated as before on our panel of white box attacks and image corruptions. Three variants targeted the GFB: one sampling the Gabor parameters from uniform distributions instead of those found in primate V1, another without high spatial frequency (SF) filters (*f* < 2cpd), and another without low SF filters (*f* > 2cpd). Two additional variants targeted the nonlinearities: one without simple cells, and another without complex cells. Finally, the sixth variant had the V1 neuronal stochasticity generator removed. Even though all VOneBlock variants are poorer approximations of primate V1, all resulting VOneResNet50 variants still had improved perturbation accuracy when compared to the base ResNet50 model (Fig. 3 A). On the other hand, all variants except that with uniformly sampled Gabor parameters showed significant deficits in robustness compared to the unmodified VOneResNet50 (Fig. 3 A, drops in perturbation accuracy between 1% and 15%).

**Figure 3:**
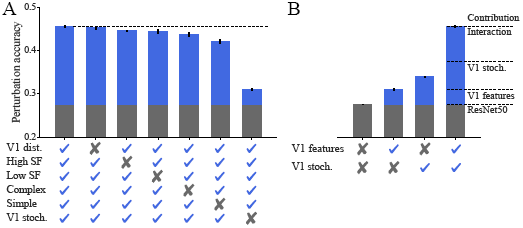
All components of the VOneBlock work in synergy to improve robustness to image perturbations. **A** Perturbation mean accuracy (top-1) for VOneResNet50 and several variations with a component of the VOneBlock removed or altered. From left to right: unmodified VOneResNet50, model with Gabor parameters sampled uniformly (within biological ranges), model without high SF Gabors, model without low SF Gabors, model without complex cells, model without simple cells, model without V1 stochasticity. Gray bars show the performance of ResNet50. Blue bars show the improvements due to the VOneBlock. Dashed line indicates the accuracy of the unmodified model. Error-bars represent SD (n=3 seeds). **B** Same as in **A** but comparing ResNet50, VOneResNet50 without V1 stochasticity, ResNet50 with V1 stochasticity added after the first block, and VOneRes-Net50. Adding V1 stochasticity to ResNet50 accounts for less than half of the total improvement of VOneResNet50, demonstrating a supralinear interaction between the V1 features and V1 stochasticity.

Interestingly, we observed that some of these changes had a highly specific effect on the type of perturbation robustness affected (Table 2 and Supplementary Section D). Removing high SF Gabors negatively affected both white box and clean accuracy while actually improving robustness to common image corruptions, particularly those of the noise and blur categories (Supplementary Table D.1). Removing complex cells only impaired white box robustness, as opposed to removing simple cells, which was particularly detrimental to performance on image corruptions. Finally, removing V1 stochasticity considerably decreased white box accuracy while improving accuracy for both clean and corrupted images.

**Table 2:** Removal of some VOneBlock components causes impairments with high specificity. Difference in accuracy (top-1) relative to the unmodified VOneResNet50, for each of the variants with removed components on overall mean, white box attacks, common corruptions, and clean images. Values are reported as mean and SD (n=3 seeds).

| Component removed | Overall | Perturbation |  |  |
| --- | --- | --- | --- | --- |
| | Mean [$\Delta\%$] | White box [$\Delta\%$] | Corruption [$\Delta\%$] | Clean [$\Delta\%$] |
| V1 dist. | -0.4±0.3 | 0.4±0.8 | -1.1±0.3 | -0.5±0.3 |
| High SF | -2.2±0.2 | -3.8±0.3 | 1.9±0.4 | **-4.7**±0.2 |
| Low SF | -0.7±0.5 | -1.6±1.1 | -0.7±0.1 | 0.1±0.3 |
| Complex | -1.0±0.6 | -4.5±1.1 | 0.6±0.4 | 0.8±0.3 |
| Simple | -3.0±0.5 | -2.0±1.1 | **-4.8**±0.5 | -2.1±0.3 |
| V1 stoch. | **-8.8**±0.3 | **-30.6**±0.4 | 1.5±0.5 | 2.6±0.1 |

The VOneBlock variant without V1 stochasticity suffered the most dramatic loss in robustness. This is not altogether surprising, as several approaches to adversarial robustness have focused on noise as a defense [15, 19, 66]. To investigate whether V1 stochasticity alone accounted for the majority of the robustness gains, we evaluated the perturbation accuracy of a ResNet50 model with V1 stochasticity added at the output of its first block. Neuronal stochasticity was implemented exactly the same way as in the VOneBlock by first applying an affine transformation to scale the activations so that they match primate V1 neuronal activity. Like the VOneResNet50, this model had stochasticity during training and inference, and showed a considerable improvement in robustness compared to the standard ResNet50. However, this improvement accounted for only a fraction of the total gains of the VOneResNet50 model (Fig. 3 B), demonstrating that there is a substantial supralinear interaction between the V1 features and the neuronal stochasticity. Merely adding V1 stochasticity to the first block of a standard CNN model does not increase robustness to the same degree as the full VOneBlock—the presence of V1 features more than doubles the contribution to perturbation robustness brought by the addition of neuronal stochasticity.

Finally, we sought to determine whether stochasticity during inference is key to defending against attacks. Thus, we analyzed the white box adversarial performance of VOneResNet50 while quenching stochasticity during the adversarial attack (Fig. 4). Remarkably, the majority of improvements in adversarial robustness originate from the neuronal stochasticity during training, indicating that V1 stochasticity induces the downstream layers to learn representations that are more robust to adversarial attacks. This is particularly interesting when contrasted with the ANT^3×3^+SIN defense, which has noise added to input images during training, but does not learn representations with notably higher robustness to white box adversarial attacks (Table 1 and Supplementary Fig. B.1).

**Figure 4:**
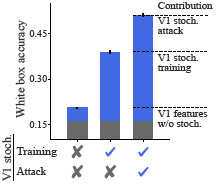
V1 stochasticity during training leads to more robust learned representations. White box accuracy (top-1) for VOneResNet50 variant without V1 stochasticity, VOneResNet50 with stochasticity removed during the white box attack and inference, and VOneResNet50. VOneResNet50 without stochasticity during attack/inference maintained over half of the robustness gains. Therefore, V1 stochasticity improves model robustness in two ways: rendering the attack itself less effective and inducing more robust representations downstream of the VOneBlock, the latter accounting for the majority of the gains on our benchmark. Gray bars show the performance of the standard ResNet50. Blue bars show the improvements due to the VOneBlock. Error-bars represent SD (n=3 seeds).

## 5 Discussion

In this work, we demonstrate that adversarial robustness in CNNs is correlated with their ability to explain primate V1 neuronal responses, and that simulating the image processing of primate V1 at the front of standard neural network architectures significantly improves their robustness to image perturbations. Notably, this approach outperformed state-of-the-art defense methods on a large benchmark consisting of adversarial attacks and common image corruptions. Despite not being constructed to this end, any component of the model front-end we removed or modified to be less like primate V1 resulted in less overall robustness, and revealed that different components improve robustness to different perturbation types. Remarkably, we find that simulated V1 stochasticity interacts synergistically with the V1 model features, and drives the downstream layers to learn representations more robust to adversarial perturbations.

Our approach bears some similarity to the pioneering study by Li et. al. [40], in which a model’s representations were regularized to approximate mouse V1, increasing its robustness in the grey-scale CIFAR dataset; however, here we go further in several key ways. First, while the mouse is gaining traction as a model for studying vision, a vast literature has established macaque vision as a quantitatively accurate model of human vision in general and human object recognition in particular [85, 86]. Visual acuity in mice is much lower than in macaques and humans [87, 79], suggesting that vision in the mouse may serve different behavioral functions than in primates. Further, the regularization approach employed by Li et. al. does not allow a clear disambiguation of which aspects of mouse V1 contribute to the improved robustness. Since the components of the VOneBlock proposed here are mappable to the brain, we can dissect the contributions of different neuronal populations in primate V1 to robustness against specific image perturbations. Finally, extending the robustness gains of biologically-constrained CNNs from gray-scale CIFAR to the full ImageNet dataset is a critical step towards real-world, human-level applications.

The gains achieved by VOneNets are substantial, particularly against white box attacks, and have tangible benefits over other defense methods. Though adversarial training still provides the strongest defense for the attack statistics it is trained on, it has significant downsides. Beyond its considerable additional computational cost during training, adversarially trained networks have significantly lower performance on clean images, corrupted images, and images perturbed with attack statistics not seen during training, implying that adversarial training in its current form may not be viable as a general defense method. In contrast, by deploying an architectural change, **VOneNets improve robustness to all adversarial attacks tested and many common image corruptions, and they accomplish this with no additional training overhead**. This also suggests that the architectural gains of the VOneNet could be stacked with other training based defense methods to achieve even greater overall robustness gains.

Relative to current methods, the success of this approach derives from engineering in a better approximation of the architecture of the most well studied primate visual area, combined with task optimization of the remaining free parameters of the downstream architecture [26]. This points to two potentially synergistic avenues for further gains: an even more neurobiologically precise model of V1 (i.e. a better VOneBlock), and an even more neurobiologically precise model of the downstream architecture. For example, one could extend the biological fidelity of the VOneBlock, in the hope that it confers even greater robustness, by including properties such as divisive normalization [48, 49] and contextual modulation [88], to name a few. In addition to V1, the retina and the Lateral Geniculate Nucleus (LGN) also play important roles in pre-processing visual information, only partially captured by the current V1 model, suggesting that extending the work done here to a retina/LGN front-end has potential to better align CNNs with human visual object recognition [89, 90]. In addition, though our initial experiments show that multiple components of our V1 model work together to produce greater robustness, the nature of the relationship between adversarial robustness and explained variance in V1 responses is far from resolved. In particular, as the VOneNet with and without stochasticity achieve very similar V1 explained variances but have markedly different levels of adversarial robustness, it is clear that much theoretical work remains in better understanding both when and why matching biology leads to more robust computer vision models.

While neuroscience has recently seen a huge influx of new neurally-mechanistic models and tools drawn from machine learning [91, 92], the most recent advances in machine learning and computer vision have been driven mostly by the widespread availability of computational resources and very large labeled datasets [93], rather than by an understanding of the relevant brain mechanisms. Under the belief that biological intelligence still has a lot of untapped potential to contribute, a number of researchers have been pushing for more neuroscience-inspired machine learning algorithms [37, 42, 32]. The work presented here shows that this aspiration can become reality—the models presented here, drawn directly from primate neurobiology, indeed require less training to achieve more human-like behavior. This is one turn of a new virtuous circle, wherein neuroscience and artificial intelligence each feed into and reinforce the understanding and ability of the other.

## Broader Impact

From a technological perspective, the ethical implications of our work are largely aligned with those of computer vision in general. While there is undoubtedly potential for malicious and abusive uses of computer vision, particularly in the form of discrimination or invasion of privacy, we believe that our work will aid in the production of more robust and intuitive behavior of computer vision algorithms. As CNNs are deployed in real-world situations, it is critical that they behave with the same level of stability as their human counterparts. In particular, they should at the very least not be confused by changes in input statistics that do not confuse humans. We believe that this work will help to bridge that gap. Furthermore, while algorithms are often thought to be impartial or unbiased, much research has shown that data driven models like current CNNs are often even more biased than humans, implicitly keying in on and amplifying stereotypes. For this reason, making new CNNs that behave more like humans may actually reduce, or at least make more intuitive, their implicit biases. Unfortunately, we note that even with our work, these issues are not resolved, yet. While we developed a more neurobiologically-constrained algorithm, it comes nowhere close to human-like behaviour in the wide range of circumstances experienced in the real world. Finally, from the perspective of neuroscience, we think that this work introduces a more accurate model of the primate visual system. Ultimately, better models contribute to a better mechanistic understanding of how the brain works, and how to intervene in the case of illness or disease states. We think that our model contributes a stronger foundation for understanding the brain and building novel medical technology such as neural implants for restoring vision in people with impairments.

## Acknowledgments and Disclosure of Funding

We thank J. Anthony Movshon and Corey M. Ziemba for access to the V1 neural dataset, MIT-IBM Satori for providing the compute necessary for this project, John Cohn for technical support, Sijia Liu and Wieland Brendel for advice on adversarial attacks, and Adam Marblestone for insightful discussion. This work was supported by the PhRMA Foundation Postdoctoral Fellowship in Informatics (T.M), the Semiconductor Research Corporation (SRC) and DARPA (J.D., M.S., J.J.D.), the Massachusetts Institute of Technology Shoemaker Fellowship (M.S.), Office of Naval Research grant MURI-114407 (J.J.D.), the Simons Foundation grant SCGB-542965 (J.J.D.), the MIT-IBM Watson AI Lab grant W1771646 (J.J.D.).

## Supplementary Material

### A Neural Explained Variance

We evaluated how well responses to given images in candidate CNNs explain responses of single V1 neurons to the same images using a standard neural predictivity methodology based on partial least square (PLS) regression [31, 32]. We used a neural dataset obtained by extracellular recordings from 102 single-units while presenting 450 different images for 100ms, 20 times each, spanning 4deg of the visual space. Results from this dataset were originally published in a study comparing V1 and V2 responses to naturalistic textures [77]. To avoid over-fitting, for each CNN, we first chose its best V1 layer on a subset of the dataset (n=135 images) and reported the final neural explained variance calculated for the chosen layer on the remaining images of the dataset (n=315 images). Model units were mapped to each V1 neuron linearly using a PLS regression model with 25 components. The mapping was performed for each neuron using 90% of the image responses in the respective dataset and tested on the remaining held-out 10% in a 10-fold cross-validation strategy. We reported neural explained variance values normalized by the neuronal internal consistency. We calculated V1 neural explained variance for a large pool of CNNs (n=30 models).

When using CNNs in a computer vision task, such as object recognition in ImageNet, their field-of-view (in degrees) is not defined since the only relevant input property is the image resolution (in pixels). However, when using a CNN to model the ventral stream, the visual spatial extent of the model’s input is of key importance to ensure that it is correctly mapped to the data it is trying to explain. We assigned a spatial extent of 8deg to all models since this value has been previously used in studies benchmarking CNNs as models of the primate ventral stream [31, 94], and is consistent with the results in Cadena et. al. [30].

### B Image Perturbations

#### B.1 White Box Adversarial Attacks

For performing white box adversarial attacks, we used untargeted projected gradient descent (PGD) [17] (also referred to as the Basic Iterative Method [95]) with *L*_∞_, *L*_2_, and *L*_1_ norm constraints. Given an image *x*, This method uses the gradient of the loss to iteratively construct an adversarial image *x_adv_* which maximizes the model loss within an *L_p_* bound around *x*. Formally, in the case of an *L*_∞_ constraint, PGD iteratively computes *x_adv_* as

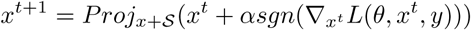

where *x* is the original image, and the *Proj* operator ensures the final computed adversarial image *x_adv_* is constrained to the space *x* + *𝓢*, here the *L*_∞_ ball around *x*. In the case of *L*_1_ or *L*_2_ constraints, at each iteration 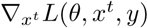 is scaled to have an *L*_1_ or *L*_2_ norm of *α*, and the *Proj* operator ensures the final *x_adv_* is within an *L*_1_ or *L*_2_ norm ball around *x*.

For our white box benchmarks, we used ‖*δ*‖_∞_ ∈ [1/1020, 1/255], ‖*δ*‖_2_ ∈ [0.15, 0.6], and ‖*δ*‖_1_ ∈ [40, 160] constraints where **δ** = *x*−*x_adv_*, for a total of six different attack variants, with two strengths under each norm constraint. We arrived at these strengths by calibrating the lower strength attack for each norm to give a reasonable degree of variance in normally trained CNNs. Because normally trained CNNs are extremely sensitive to adversarial perturbations, this resulted in perturbations well below the level of perceptibility, and in most cases, particularly with the *L*_∞_ constraint, perturbations that would not even be rendered differently by a monitor for display. We set the high attack strength at 4 times the low value, which brought standard ImageNet trained CNNs to nearly chance performance. Still, even this higher perturbation strength remained practically imperceptible (Fig. B.3 left). Each perturbation was computed with 64 PGD iterations and a step size *α* = *ϵ*/32, where *ϵ* = ‖*δ*‖_*p*_, for the same 5000 images from the ImageNet validation set, on a per model basis, and final top-1 accuracy was reported. The Adversarial Robustness Toolkit [96] was used for computing the attacks.

While adversarial formulations using Lagrangian relaxation like the C&W attack [7] and the EAD attack [8] are known to be more effective than PGD for finding minimal perturbations under *L*_2_ and *L*_1_ constraints, these formulations involve computationally expensive line searches, often requiring thousands of iterations to converge, making them computationally difficult to scale to 5000 ImageNet samples for a large number of models. Furthermore, because some of the models we tested operate with stochasticity during inference, we found that other state-of-the-art attacks formulated to efficiently find minimal perturbation distances [9, 10] were generally difficult to tune, as their success becomes stochastic as they near the decision boundary. Thus we proceeded with the PGD formulation as we found it to be the most reliable, computationally tractable, and conceptually simple to follow. We share our model and weights, and encourage anyone interested to attack our model to verify our results.

Although the stochastic models were implemented with the reparameterization trick for random sampling, which leaves the gradients intact [83], we performed several sanity checks to make sure that the white box attacks were effectively implemented. Following the advice of Athalye et. al. [68] and Carlini et. al. [84], we observed that in all of the white box attacks, increasing the attack strength decreased model performance, eventually bringing accuracy to zero (Fig. B.1), and additionally that for any given constraint and attack strength, increasing PGD iterations from one to many increased attack effectiveness (Fig. B.2). Furthermore, to be extra cautious, we followed the method Athalye et. al. [68] used to break Stochastic Activation Pruning [19] and replaced ▽*_x_f*(*x*) with 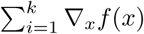, effectively performing Monte Carlo (MC) sampling of the loss gradient at every PGD iteration. While with high perturbation strength even *k* = 1 was sufficient to bring our model’s accuracy to zero, we find that *k* = 10 generally achieved higher attack success at intermediate and large perturbation strengths (Table B.1), and thus we used this approach for all attacks on stochastic models.

**Figure B.1:**
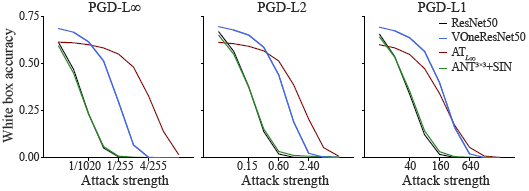
VOneResNet50 improves robustness to white box attacks in a wide range of perturbation strengths. White box accuracy perturbation strength curves for PGD attacks with constraints *L*_∞_, *L*_2_ and *L*_1_ for ResNet50, VOneResNet50, AT_*L*∞_, and ANT^3×3^+SIN. Adding the VOneBlock to ResNet50 consistently improves robustness under all constraints and attack strength—VOneResNet50 can withstand perturbations roughly 4 times higher than the standard ResNet50 for the same performance level. Remarkably, VOneResNet50 outperformed the adversarially trained model (AT_*L*∞_) for a considerable range of perturbation strengths, particularly with the *L*_1_ constraint. Adversarial noise training combined with Stylized ImageNet (ANT^3×3^+SIN) has virtually no improvement in robustness to white box attacks. In all white box attacks and for all models, increasing the attack strength decreased performance, eventually bringing accuracy to zero.

**Figure B.2:**
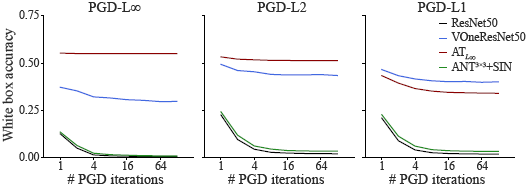
White box attack effectiveness converges for large number of PGD iteration steps. White box accuracy iteration curves for PGD attacks with ‖*δ*‖_∞_ = 1/255, ‖*δ*‖_2_ = 0.6, ‖*δ*‖_1_ = 160 constraints. Increasing the number of PGD iteration steps makes the attack more effective, generally converging after 32 iterations.

**Table B.1:** Improving the effectiveness of white box attacks with Monte Carlo sampling. White box attack accuracy (top-1) under different norm constraints and white box overall mean using standard PGD and PGD with Monte Carlo sampling of the loss gradients (PGD-MC, *k* = 10) on VOneResNet50 for 64 PGD iteration steps. For all stochastic models we used PGD with Monte Carlo sampling in the white box attacks throughout the paper. Values are mean and SD (n=3 seeds).

| Attack type | White box | PGD-$L_1$ | | PGD-$L_2$ | | PGD-$L_\infty$ | |
| --- | --- | --- | --- | --- | --- | --- | --- |
|  | Mean<br>[%] | 40<br>[%] | 160<br>[%] | 0.15<br>[%] | 0.60<br>[%] | 1/1020<br>[%] | 1/255<br>[%] |
| Standard PGD | $52.8 \pm 0.2$ | $64.4 \pm 0.4$ | $42.2 \pm 0.1$ | $65.2 \pm 0.3$ | $45.3 \pm 0.4$ | $63.4 \pm 0.4$ | $36.4 \pm 0.4$ |
| PGD-MC ($k=10$) | $51.1 \pm 0.4$ | $64.2 \pm 0.5$ | $40.4 \pm 0.5$ | $65.3 \pm 0.4$ | $44.3 \pm 0.3$ | $61.9 \pm 0.4$ | $30.3 \pm 0.5$ |

Although VOneResNet50 was not designed to explicitly improve robustness to white box attacks, it offers substantial gains over the standard ResNet50 model (Table 1). A closer inspection of the white box accuracy perturbation strength curves (Fig. B.1) reveals that VOneResNet50 can withstand perturbations roughly 4 times higher than the standard ResNet50 for the same performance level (blue curve is shifted to the right of the black curve by a factor of 4). To help visualize how this difference translates into actual images, we show in Fig. B.3 examples of white box adversarial images for both ResNet50 and VOneResNe50 for the perturbation strengths that bring the respective models to nearly chance performance.

**Figure B.3:**
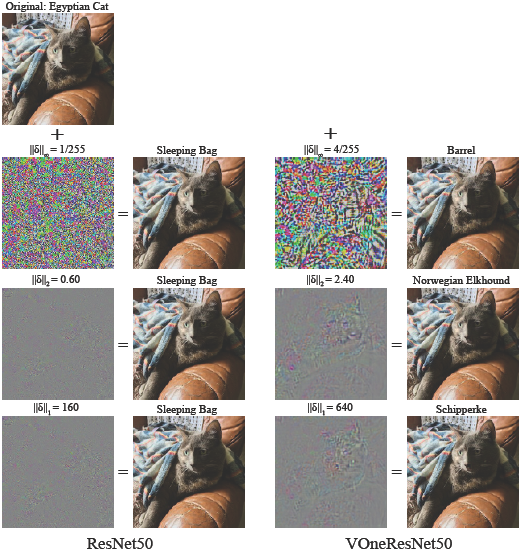
Comparison of adversarial images for ResNet50 and VOneResNet50. Visualization of white box attacks under the 3 different norm constraints (*L*_∞_, *L*_2_, and *L*_1_) for the perturbation strength that brings the model to nearly chance performance (Fig. B.1). Left, ResNet50; right, VOneResNet50. For each attack, the perturbation is shown on the left, scaled for visualization, and the perturbed image appears next to it. Perturbation strengths and classes of the adversarial images are shown over the images.

#### B.2 Additional attack controls

The field of adversarial robustness is littered with examples of proposed defense methods that were quickly defeated with some minor attack optimizations or adaptations [68, 84, 97]. For this reason, we took great care to verify our claims of improved robustness of the VOneNets. In addition to the implementation details and controls described in the previous section, we validated our results by further optimizing our attacks on both the standard stochastic VOneResNet50 and the VOneResNet50 with stochasticity removed during the attack and inference.

First, on the standard stochastic VOneResNet50, we ensured that our attack hyper-parameters, including step-size (*α*), number of gradient samples (*k*), and PGD iterations, are sufficient for evaluating the network’s robustness. For PGD with ‖*δ*‖_∞_ = 1/255 with 64 iterations, we performed a large grid search over *α* and *k*, observing only a marginal increase in the attack effectiveness (Fig. B.4). For the most effective step size (*α* = *ϵ*/8), increasing the number of gradient samples beyond *k* = 16 no longer decreases accuracy under the attack. At *k* = 128, VOneResNet50 accuracy is only reduced from 29.15% to 26.0%, remaining a large margin above ResNet50 accuracy of 0.8%. Additionally, increasing the number of PGD iterations all the way up to 1000 (with *α* = *ϵ*/8 and *k* = 16), only further reduced performance by 0.5%. Finally, using 5-10 random restarts did not improve the attack beyond the most effective *k* and *α* settings.

**Figure B.4:**
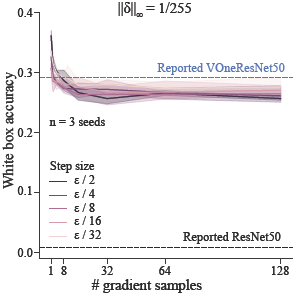
Further attack optimization does not overturn results. White box accuracy for PGD with ‖*δ*‖_∞_ = 1/255, 64 iterations on 500 images. Large grid search over step size and number of gradient samples only marginally improves attack effectiveness. Further attack optimization only marginally reduces model performance. Lines represent mean and shaded areas SD (n=3 seeds).

Second, while boundary attacks are not ideal against the standard stochastic VOneResNet50 due to the shifting boundary line, we investigated VOneResNet50 without stochasticity during attack/inference using the Brendel and Bethge (B&B) attack [10], a powerful white box adversary that aims to find minimal adversarial perturbations along the decision boundary of the network. We reasoned that if the B&B attack offers similar results to those observed using PGD, this is a good sanity check that our observed robustness is reliable, and not a result of masked gradients around the input or a quirk of PGD. Using the FoolBox [98] implementation of the *L*_∞_ B&B attack initialized with 500 successful adversarial images generated using PGD ‖*δ*‖_∞_ = 8/255, we see nearly identical scores as compared to standard PGD with optimal hyper-parameters ‖*δ*‖_∞_ = 1/1020 and PGD ‖*δ*‖_∞_ = 1/255.

We conclude that these additional controls do not qualitatively change our reported results and provide further evidence that the observed improved robustness of the VOneNets is real and not an artifact.

#### B.3 Common Image Corruptions

For evaluating model robustness to common image corruptions, we used ImageNet-C, a dataset publicly available at https://github.com/hendrycks/robustness [13]. It consists of 15 different types of corruptions, each at five levels of severity for a total of 75 different perturbations (Fig. B.5). The corruption types fall into four categories: noise, blur, weather, and digital effects. Noise includes Gaussian noise, shot noise, and impulse noise; blur includes motion blur, defocus blur, zoom blur, and glass blur; weather includes snow, frost, fog, and brightness; digital effects includes JPEG compression, elastic transformations, contrast, and pixelation. For every corruption type and at every severity, we tested on all 50,000 images from the ImageNet validation set. We note that these images have been JPEG compressed by the original authors for sharing, which is known to have minor but noticeable effect on final network performance.

**Figure B.5:**
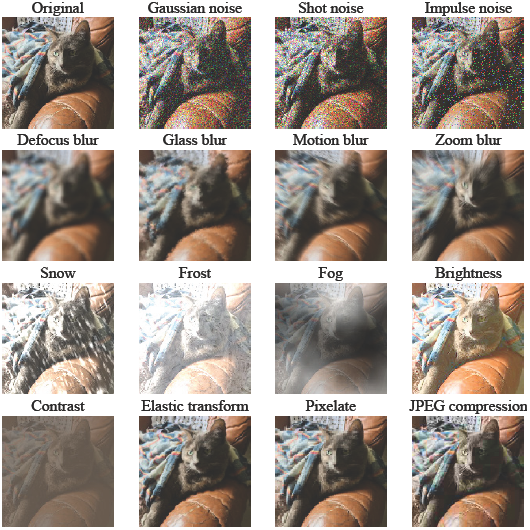
Common image corruptions with intermediate severity. Visualization of all 15 common image corruption types evaluated at severity = 3. First row, original image, followed by the noise corruptions; second row, blur corruptions; third row, weather corruptions; fourth row, digital corruptions.

**Figure B.6:**
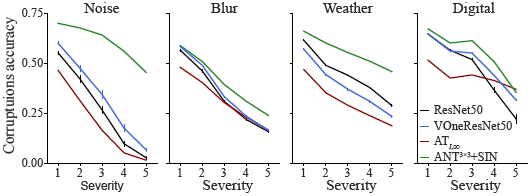
VOneResNet50 improves robustness to noise, blur and digital corruptions. Corruption accuracy severity curves for the four categories of common image corruptions: noise, blur, weather, and digital. Adding the VOneBlock to ResNet50 improves robustness to all corruption categories except weather. The adversarially trained model (AT_*L*∞_) is considerably worse at common image corruptions. Adversarial noise combined with Stylized ImageNet training (ANT^3×3^+SIN) is consistently more robust to all corruption categories. For ResNet50 and VOneResNet50, error-bars represent SD (n=3 seeds).

**Table B.2:** Detailed common image corruption results. Adding the VOneBlock to ResNet50 improves robustness to several of the corruption types. However, for others, particularly fog and contrast, robustness decreases. Interestingly, these are the corruption types that the adversarially trained model (AT_*L*∞_) also does the worst. Adversarial noise combined with Stylized ImageNet training (ANT^3×3^+SIN) is the most robust model for most of the corruption types. For ResNet50 and VOneResNet50, values are mean and SD (n=3 seeds).

| Model | Noise |  |  | Blur |  |  |  |
| --- | --- | --- | --- | --- | --- | --- | --- |
|  | Gaussian [%] | Shot [%] | Impulse [%] | Defocus [%] | Glass [%] | Motion [%] | Zoom [%] |
| ResNet50 [4] | 29.6±1.1 | 27.9±0.9 | 24.5±1.2 | 38.8±0.7 | 26.1±0.6 | 37.9±0.4 | 34.5±0.7 |
| $AT_{L_{\infty}}$ [76] | 23.6 | 22.3 | 14.8 | 24.4 | 34.3 | 33.2 | 34.9 |
| $ANT^{3 \times 3} + SIN$ [59] | **61.6** | **60.7** | **59.9** | **41.5** | **38.6** | **46.0** | **37.3** |
| VOneResNet50 | 34.6±1.4 | 33.4±1.2 | 31.9±1.5 | 37.8±0.1 | 35.7±0.1 | 37.4±0.2 | 34.0±0.2 |

| Model | Weather |  |  |  | Digital |  |  |  |
| --- | --- | --- | --- | --- | --- | --- | --- | --- |
|  | Snow [%] | Frost [%] | Fog [%] | Bright. [%] | Contrast [%] | Elastic [%] | Pixelate [%] | JPEG [%] |
| ResNet50 | 30.1±0.5 | 36.7±0.4 | 43.6±0.8 | 66.8±0.2 | 38.8±0.5 | 44.8±0.3 | 47.0±2.7 | 55.1±1.0 |
| $AT_{L_{\infty}}$ | 29.5 | 31.2 | 8.4 | 54.1 | 10.2 | 48.4 | 56.3 | 58.7 |
| $ANT^{3 \times 3} + SIN$ | **47.7** | **52.2** | **54.9** | **68.2** | **49.3** | **52.7** | 58.9 | 59.3 |
| VOneResNet50 | 25.2±0.5 | 36.8±0.4 | 30.1±0.5 | 62.4±0.3 | 28.5±0.5 | 48.7±0.1 | **63.3±0.2** | **61.0±0.3** |

#### B.4 Other defense methods

We compared the performance of VOneResNet50 in the described computer vision benchmarks with two training-based defense methods using the ResNet50 architecture: adversarial training with a ‖*δ*‖_∞_ = 4/255 constraint (AT_*L*∞_) [76], and adversarial noise with Stylized ImageNet training (ANT^3×3^+SIN) [59].

The AT_*L*∞_ model was downloaded from the publicly available adversarially trained models at https://github.com/MadryLab/robustness [76]. This model was trained following the adversarial training paradigm of Madry et. al. [17], to solve the min-max problem,

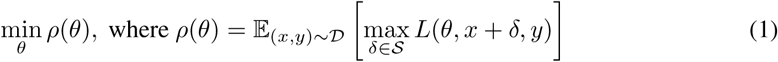

Here, the goal is to learn parameters *θ* minimizing the loss *L* for training images *x* and labels *y* drawn from *D*, while perturbing *x* with **δ** ∈ *𝓢* to maximally increase the loss. In this case, *𝓓* is ImageNet, and PGD with ‖*δ*‖_∞_ = 4/255 is used to find the perturbation maximizing the loss for a given (*x, y*) pair. The ‖*δ*‖_∞_ = 4/255 constraint model was selected to compare against because it had the best performance on our conglomerate benchmark of adversarial attacks and common image corruptions.

The ANT^3×3^+SIN model was obtained from https://github.com/bethgelab/game-of-noise. Recently, Rusak et. al. showed that training a ResNet50 model with several types of additive input noise improved the robustness to common image corruptions [59]. Inspired by this observation, the authors decided to train a model while optimizing a noise distribution that maximally confuses it. Similarly to standard adversarial training, this results in solving a min-max problem,

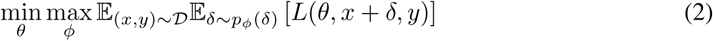

where *p*_*φ*_ (*δ*) is the maximally confusing noise distribution. The main difference to regular adversarial training in Equation 1, is that while in the former *δ* is optimized directly, here *δ* is found by optimizing a constrained distribution with local spatial correlations. Complementing the adversarial noise training with Stylized ImageNet training [25] produced the model with the current best robustness in the ImageNet-C benchmark as a standalone method.

### C VOneNet implementation details

#### C.1 Convolutional layer

The convolutional layer of the VOneBlock is a mathematically parameterized Gabor Filter Bank (GFB). We set the stride of the GFB to be four, originating a 56×56 spatial map of activations. Since the number of channels in most CNNs’ first convolution is relatively small (64 in the architectures adapted), we used a larger number in the VOneBlock so that the Gabors would cover the large parameter space and better approximate primate V1. We set the main VOneNet models to contain 512 channels equally split between simple and complex cells (see Fig. C.2 A for an analysis of how the number of channels affects performance). Each channel in the GFB convolves a single color channel from the input image.

The Gabor function consists of a two-dimensional grating with a Gaussian envelope and is described by the following equation:

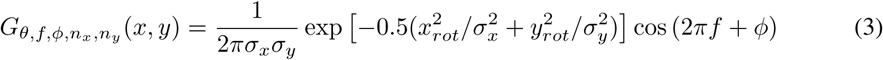

where

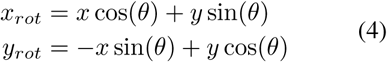

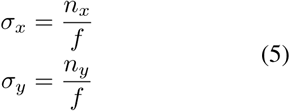

*x_rot_* and *y_rot_* are the orthogonal and parallel orientations relative to the grating, *θ* is the angle of the grating orientation, *f* is the spatial frequency of the grating, *ϕ* is the phase of the grating relative to the Gaussian envelope, and *σ_x_* and *σ_y_* are the standard deviations of the Gaussian envelope orthogonal and parallel to the grating, which can be defined as multiples (*n_x_* and *n_y_*) of the grating cycle (inverse of the frequency).

Although the Gabor filter greatly reduces the number of parameters defining the linear spatial component of V1 neurons, it still has five parameters per channel. Fortunately, there is a vast literature in neurophysiology with detailed characterizations of primate V1 response properties which can be used to constrain these parameters. To instantiate a VOneBlock with *C*_*V*1_ channels, we sampled *C*_*V*1_ values for each of the parameters according to an empirically constrained distribution (Table C.1). Due to the resolution of the input images, we limited the ranges of spatial frequencies (*f* < 5.6cpd) and number of cycles (*n* > 0.1).

**Table C.1:** Empirical distributions used to sample the GFB parameters in the VOneBlock.

| Parameter | Reference |  |
| --- | --- | --- |
| $\theta$ | De Valois et. at. [78] | Fig. 5: Orientation tuning (foveal) |
| $f$ | De Valois et. at. [79] | Fig. 6: Peak spatial freq. (X cells foveal) |
| $\phi$ | - | Uniform distribution $\in [0, 360]$ deg |
| $n_x, n_y$ | Ringach 2002 [80] | Fig. 4: $(n_x, n_y)$ joint distribution |

Critical to the correct implementation of the biologically-constrained parameters of the GFB is the choice of the model’s field of view in degrees. As previously mentioned in Section A, we used 8deg as the input spatial extent for all CNN models. It’s important to note that the set of spatial filters in the VOneBlock’s GFB differs considerably from those in the first convolution in most CNNs (Fig. C.1). While standard CNNs learn filters that resemble Gabors in their input layer [1, 99, 100], due the limited span of their kernels, they do not vary significantly in size and spatial frequency. V1 neurons, on the other hand, are known to exhibit a wide range of receptive field properties. This phenomena is captured in the VOneBlock with the spatial frequencies and receptive field sizes of Gabors ranging more than one order of magnitude. Due to this high variability, we set the convolution kernel to be 25 × 25px, which is considerably larger than those in standard CNNs (Fig. C.1).

**Figure C.1:**
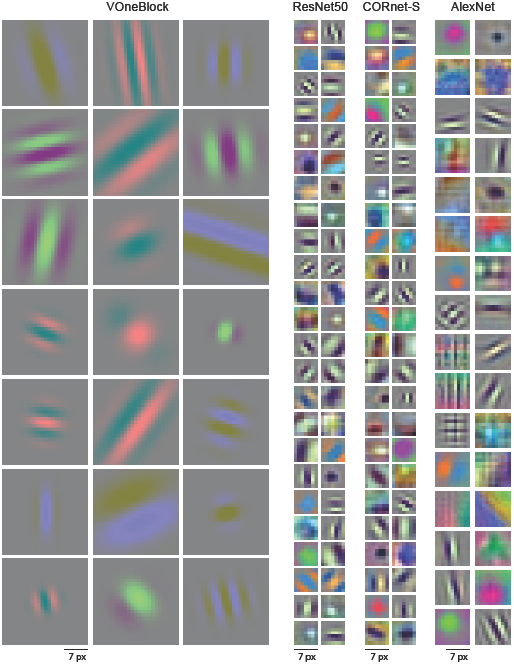
Comparison between filters in the first convolution of standard CNNs and the VOneBlock. Example filters from the first convolution in the VOneBlock, ResNet50, CORnet-S, and AlexNet (from left to right). VOneBlock filters are all parameterized as Gabors, varying considerably in size and spatial frequency. Standard CNNs have some filters with shapes other than Gabors, particularly center-surround, and are limited in size by their small kernel. Kernel sizes are 25px, 7px, 7px, and 11px for VOneBlock, ResNet50, CORnet-S, and AlexNet, respectively.

#### C.2 Nonlinear layer

VOneBlock’s nonlinear layer has two different nonlinearities that are applied to each channel depending on its cell type: a rectified linear transformation for simple cells (6), and the spectral power of a quadrature phase-pair (7) for complex cells:

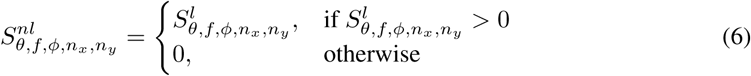

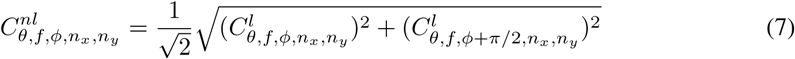

where 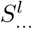 and 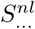 are the linear and nonlinear responses of a simple neuron and 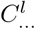 and 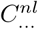 are the same for a complex neuron.

#### C.3 Neuronal stochasticity generator

In awake monkeys, spike trains of V1 neurons are approximately Poisson, i.e. the variance and mean of spike counts, in a given time-window, over a set of repetitions are roughly the same [82]. We incorporated stochasticity into the VOneBlock to emulate this property of neuronal responses. Since the Poisson distribution is not continuous, it breaks the gradients in a white box attack giving a false sense of robustness [68]. In order to avoid this situation and facilitate the evaluation of the model’s real robustness, our neuronal stochasticity generator as implemented uses a continuous, second-order approximation of Poisson noise by adding Gaussian noise with variance equal to the activation:

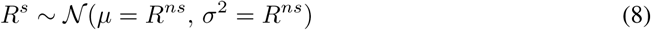

where *R^ns^* and *R^s^* are the non-stochastic and stochastic responses of a neuron.

To approximate the same levels of neuronal stochasticity of primate V1 neurons, it is critical that the VOneBlock activations are on the same range as the V1 neuronal responses (number of spike counts in a given time-window). Thus, we applied an affine transformation to the activations so that both the mean stimulus response and the mean baseline activity are the same as those of a population of V1 neurons (Table C.2 shows the mean responses and spontaneous activity of V1 neurons measured in different time-windows).

**Table C.2:** Mean stimulus responses and spontaneous activity of a population of V1 neurons for different time-windows.

| Time-window<br>[ms] | Stimulus response<br>[spikes] | Spontaneous activity<br>[spikes] |
| --- | --- | --- |
| 25 | 0.324 | 0.073 |
| 50 | 0.655 | 0.145 |
| 100 | 1.05 | 0.29 |
| 200 | 1.38 | 0.58 |

Since the exact time-window that responses in V1 are integrated during visual perception is still an open question, we considered a time-window of 25ms (Fig. C.2 B). In order to keep the outputs of the VOneBlock on a range that does not deviate considerably from the typical range of activations in CNNs, so that the model can be efficiently trained using standard training parameters, we applied the inverse transformation to scale the outputs back to their original range after the stochastic layer.

#### C.4 Hyper-parameters and other design choices

We developed VOneNets from three different standard CNN architectures: ResNet50, CORnet-S, and AlexNet. As previously mentioned, we replaced the first block of each architecture by the VOneBlock and the transition layer to create the respective VOneNet. Table C.3 contains the layers that were removed for each base architecture. Except CORnet-S, the layers removed only contained a single convolution, nonlinearity and maxpool. Since CORnet-S already had a pre-committed set of layers to V1, we replaced them by the VOneBlock. The torchvision implementation of AlexNet contains a combined stride of eight in the removed layers (four in the convolution and two in the maxpool), followed by a stride of one in the second convolution (outside of the removed layers). In order to more easily adapt it to a VOneBlock with a stride of four, we slightly adapted AlexNet’s architecture so that it had strides of two in these three layers (first convolution, first maxpool, and second convolution). The results shown in Fig. 2 were obtained using this modified architecture.

**Table C.3:** Layers removed from each base CNN architecture for creating the VOneNets.

| Architecture | Layers removed |
| --- | --- |
| AlexNet | Conv/ReLU/MaxPool |
| ResNet50 | Conv/Batch-Norm/ReLU/MaxPool |
| CORnet-S | Conv/Batch-Norm/ReLU/MaxPool/Conv/Batch-Norm/ReLU |

The VOneNet architecture was designed to have as few hyper-parameters as possible. When possible these were either constrained by the base CNN architecture or by neurobiological data: most parameters of the GFB were instantiated by sampling from neuronal distributions of primate V1 neurons; the kernel size was set to 25×25px to capture the high variability in Gabor sizes; like all other CNN models, the spatial extent of the field of view was set to 8deg (Section A). Nevertheless, there were two hyper-parameters where the choice was rather arbitrary: the number of channels in the VOneBlock, and the time-window for integrating V1 responses to scale the activations prior to the stochasticity generator.

Since primate V1 has neurons tuned to a wide range of spatial frequencies, for each orientation and at any given location of the visual space [101], we expected that a large number of channels would be required to cover all of the combinations of Gabor parameters. For the VOneResNet50 architecture, we varied the number of channels in the VOneBlock between 64 and 1024, equally split between simple and complex cells, and measured the performance of the several models after training (Fig. C.2 A). Clean ImageNet and common image corruption performance improved with channel number. Remarkably, the opposite happened for white box accuracy, with variants of VOneResNet50 with 64 and 128 channels achieving the highest robustness to white box attacks. This result is particularly interesting since it had been shown that increasing the number of channels in all layers of a CNN improves robustness to white box attacks in both standard and adversarially trained models [17]. Therefore, this shows that the improvement in white box robustness of VOneNets, when compared to their base models, cannot be attributed to the higher number of channels. For the main model, we set the channel number to be 512 as it offered a good compromise between the different benchmarks. Regarding the time-window for integrating V1 responses, we observed small effects on the models’ performance when varying it between 25 and 200ms. We chose 25ms for the main model due to a small trend for higher robustness to white box attacks with shorter time-windows.

**Figure C.2:**
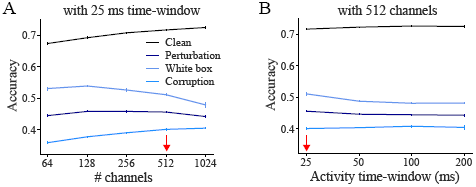
Exploration of VOneBlock’s hyper-parameters in the VOneResNet50 architecture. **A** Performance of different VOneResNet50 models with varying number of channels in the VOneBlock. **B** Same as in **A** but varying time-window for integrating neuronal activity. Error-bars show SD (n=3 seeds). Red arrows represent the values used in the main model.

#### C.5 Training details

We used PyTorch 0.4.1 and trained the model using ImageNet 2012 [34]. Images were preprocessed (1) for training—random crop to 224 × 224 pixels and random flipping left and right; (2) for validation—central crop to 224 × 224 pixels. Preprocessing was followed by normalization— subtraction and division by [0.5, 0.5, 0.5]. We used a batch size of 256 images and trained either on 2 GPUs (NVIDIA Titan X / GeForce 1080Ti) or 1 GPU (QuadroRTX6000 or V100) for 70 epochs. We use step learning rate scheduling: 0.1 starting learning rate, divided by 10 every 20 epochs. For optimization, we use Stochastic Gradient Descent with a weight decay 0.0001, momentum 0.9, and a cross-entropy loss between image labels and model predictions (logits).

Model weights and code are available at https://github.com/dicarlolab/vonenet.

### D VOneBlock variant details

To investigate which components of the VOneBlock are responsible for the increased robustness, we created six variants of the VOneBlock by removing or modifying one of its parts. In all cases, we trained from scratch a corresponding variant of VOneResNet50 with the modified VOneBlock. All other properties of the model and components of the VOneBlock were left unmodified. Here we describe in more detail each of these variants (the name of the variant refers to the component removed):

- *V1 distributions*: the GFB parameters were sampled from uniform distributions with the same domain as the empirical distributions used in the default VOneBlock (Table C.1).
- *Low SF*: when sampling the spatial frequencies for the GFB, neurons with low peak spatial frequencies (*f* < 2cpd) were removed from the empirical distribution [79].
- *High SF*: similar to the previous variant but removing neurons with high peak spatial frequencies (*f* > 2cpd).
- *Complex NL*: complex cell channels were removed from the VOneBlock and replaced by simple cell channels.
- *Simple NL*: simple cell channels were removed from the VOneBlock and replaced by complex cell channels.
- *V1 stochasticity*: V1 stochasticity generator was removed from the VOneBlock.

While all of the variants resulted in worse overall mean performance than the unmodified VOneResNet50 (Table 2), some improved specific benchmarks: the variant without high SF had higher accuracy under noise and blur corruptions (Table D.1), the variant without V1 stochasticity had higher accuracy to clean images and images with noise and weather corruptions (Tables 2 and D.1), and the variant without V1 distributions had higher accuracy to white box attacks with high *L*_∞_ perturbations (Table D.2).

**Table D.1:** Difference between the common image corruption accuracies of the several VOneResNet50 variants and the unmodified VOneResNet50. Removal of simple cells reduced accuracy to all common image corruption types. Removal of some components of the VOneBlock improved accuracy in specific common image corruption: removing high SF consistently improved performance for noise and blur corruptions, and removing V1 stochasticity improved performance for noise and weather corruptions.

| Component removed | Noise |  |  | Blur |  |  |  |
| --- | --- | --- | --- | --- | --- | --- | --- |
| | Gaussian<br>[$\Delta\%$] | Shot<br>[$\Delta\%$] | Impulse<br>[$\Delta\%$] | Defocus<br>[$\Delta\%$] | Glass<br>[$\Delta\%$] | Motion<br>[$\Delta\%$] | Zoom<br>[$\Delta\%$] |
| V1 dist. | -2.8±0.9 | -2.4±0.7 | -2.9±1.1 | -1.6±0.1 | 0.4±0.2 | -0.8±0.3 | -0.2±0.5 |
| Low SF | -1.4±0.5 | -1.0±0.2 | -1.5±0.5 | -0.6±0.6 | -1.1±0.2 | -1.2±0.4 | -0.5±0.7 |
| High SF | **6.8**±1.3 | **7.3**±1.1 | **7.0**±1.3 | **5.6**±0.3 | **11.5**±0.3 | **3.7**±0.0 | **6.1**±0.1 |
| Complex | 1.0±1.6 | 1.2±1.3 | 0.7±1.8 | 1.2±0.2 | -1.5±0.1 | 0.2±0.8 | 0.4±1.1 |
| Simple | -6.1±1.6 | -5.4±1.5 | -6.4±1.9 | -4.1±0.4 | -4.2±0.4 | -4.9±0.5 | -5.4±0.3 |
| V1 stoch. | 2.2±2.5 | 2.2±2.5 | 2.1±2.9 | 3.3±0.1 | -5.5±0.1 | 0.9±0.7 | 0.9±0.1 |

| Component removed | Weather |  |  |  | Digital |  |  |  |
| --- | --- | --- | --- | --- | --- | --- | --- | --- |
| | Snow<br>[$\Delta\%$] | Frost<br>[$\Delta\%$] | Fog<br>[$\Delta\%$] | Bright.<br>[$\Delta\%$] | Contrast<br>[$\Delta\%$] | Elastic<br>[$\Delta\%$] | Pixelate<br>[$\Delta\%$] | JPEG<br>[$\Delta\%$] |
| V1 dist. | -0.2±0.6 | -0.9±0.6 | -1.9±1.1 | -0.8±0.5 | -1.4±0.9 | 0.3±0.4 | -1.7±0.3 | **0.8**±0.5 |
| Low SF | -0.4±0.3 | -0.6±0.4 | 0.9±0.3 | -0.2±0.2 | -0.4±0.5 | -0.9±0.5 | -1.3±0.2 | -0.5±0.4 |
| High SF | 1.4±1.0 | -0.7±0.7 | -9.5±0.4 | -5.2±0.4 | -4.4±0.5 | **2.4**±0.2 | **1.9**±0.4 | -4.6±0.4 |
| Complex | 0.7±0.4 | **0.4**±0.3 | 4.3±0.2 | 1.3±0.2 | 2.9±0.2 | -0.5±0.2 | -1.2±0.0 | -1.9±0.5 |
| Simple | -5.0±0.5 | -9.0±0.5 | -4.4±0.5 | -3.9±0.2 | -4.8±0.4 | -3.9±0.5 | -3.5±0.4 | -1.8±0.2 |
| V1 stoch. | **2.5**±0.4 | **0.3**±0.3 | **12.0**±0.5 | **2.6**±0.3 | **10.0**±0.8 | -2.8±0.3 | -4.0±0.7 | -4.0±0.4 |

**Table D.2:** Difference between the white box attack accuracies of the several VOneResNet50 variants and the unmodified VOneResNet50. Removal of V1 stochasticity considerably reduced accuracy to all white box attacks. Removal of V1 distributions improved accuracy for the *L*_∞_ white box attack with high perturbation strength.

| Component removed | PGD-L1 | | PGD-L2 | | PGD-$L_\infty$ | |
| --- | --- | --- | --- | --- | --- | --- |
| | 40<br>[$\Delta\%$] | 160<br>[$\Delta\%$] | 0.15<br>[$\Delta\%$] | 0.60<br>[$\Delta\%$] | 1/1020<br>[$\Delta\%$] | 1/255<br>[$\Delta\%$] |
| V1 dist. | -0.5±0.8 | 0.1±1.1 | -0.8±0.2 | 0.6±1.0 | 0.3±0.3 | **3.0**±1.8 |
| Low SF | -0.8±1.2 | -3.1±1.1 | -0.6±1.0 | -2.3±0.8 | -0.8±1.3 | -1.8±1.0 |
| High SF | -2.2±0.9 | -4.7±2.6 | -2.6±1.5 | -5.1±1.2 | -2.6±1.4 | -5.5±1.1 |
| Complex | -1.3±0.9 | -7.4±1.6 | -1.1±0.9 | -7.2±1.2 | -1.9±1.0 | -7.9±1.2 |
| Simple | -2.5±0.8 | -1.4±1.4 | -2.9±1.1 | -1.7±1.2 | -2.7±1.2 | -0.6±0.8 |
| Noise | -23.7±0.9 | -36.5±0.8 | -21.6±0.5 | -40.1±0.4 | -32.2±0.4 | -29.2±0.4 |

## Reference

[1] Alex Krizhevsky, Ilya Sutskever, and Geoffrey E Hinton. “ImageNet Classification with Deep Convolutional Neural Networks”. In: Advances in Neural Information Processing Systems 25. Ed. by F Pereira et al. Curran Associates, Inc., 2012, pp. 1097–1105.

[2] Christian Szegedy et al. “Going Deeper with Convolutions”. In: (Sept. 2014). arXiv: 1409.4842 [cs.CV].

[3] Karen Simonyan and Andrew Zisserman. “Very Deep Convolutional Networks for Large-Scale Image Recognition”. In: (Sept. 2014). arXiv: 1409.1556 [cs.CV].

[4] Kaiming He et al. “Deep Residual Learning for Image Recognition”. In: (Dec. 2015). arXiv: 1512.03385 [cs.CV].

[5] Kaiming He et al. “Delving deep into rectifiers: Surpassing human-level performance on imagenet classification”. In: Proceedings of the IEEE International Conference on Computer Vision 2015 Inter (2015), pp. 1026–1034. ISSN: 15505499. DOI: 10.110. arXiv: 1502.01852.

[6] Christian Szegedy et al. “Intriguing properties of neural networks”. In: (Dec. 2013). arXiv: 1312.6199 [cs.CV].

[7] Nicholas Carlini and David Wagner. “Towards Evaluating the Robustness of Neural Networks”. In: (Aug. 2016). arXiv: 1608.04644 [cs.CR].

[8] Pin-Yu Chen et al. “EAD: Elastic-Net Attacks to Deep Neural Networks via Adversarial Examples”. In: (Sept. 2017). arXiv: 1709.04114 [stat.ML].

[9] Jérôme Rony et al. “Decoupling Direction and Norm for Efficient Gradient-Based L2 Adversarial Attacks and Defenses”. In: (Nov. 2018). arXiv: 1811.09600 [cs.CV].

[10] Wieland Brendel et al. “Accurate, reliable and fast robustness evaluation”. In: (July 2019). arXiv: 1907.01003 [stat.ML].

[11] Samuel Dodge and Lina Karam. “A Study and Comparison of Human and Deep Learning Recognition Performance Under Visual Distortions”. In: (May 2017). arXiv: 1705.02498 [cs.CV].

[12] Robert Geirhos et al. “Generalisation in humans and deep neural networks”. In: (Aug. 2018). arXiv: 1808.08750 [cs.CV].

[13] Dan Hendrycks and Thomas Dietterich. “Benchmarking Neural Network Robustness to Common Corruptions and Perturbations”. In: (Mar. 2019). arXiv: 1903.12261 [cs.LG].

[14] Nilaksh Das et al. “Keeping the Bad Guys Out: Protecting and Vaccinating Deep Learning with JPEG Compression”. In: (May 2017). arXiv: 1705.02900 [cs.CV].

[15] Xuanqing Liu et al. “Towards Robust Neural Networks via Random Self-ensemble”. In: (Dec. 2017). arXiv: 1712.00673 [cs.LG].

[16] Weilin Xu, David Evans, and Yanjun Qi. “Feature Squeezing: Detecting Adversarial Examples in Deep Neural Networks”. In: arXiv [cs.CV] (Apr. 2017).

[17] Aleksander Madry et al. “Towards Deep Learning Models Resistant to Adversarial Attacks”. In: (June 2017). arXiv: 1706.06083 [stat.ML].

[18] Yang Song et al. “PixelDefend: Leveraging Generative Models to Understand and Defend against Adversarial Examples”. In: (Oct. 2017). arXiv: 1710.10766 [cs.LG].

[19] Guneet S Dhillon et al. “Stochastic Activation Pruning for Robust Adversarial Defense”. In: (Mar. 2018). arXiv: 1803.01442 [cs.LG].

[20] Jacob Buckman et al. “Thermometer Encoding: One Hot Way To Resist Adversarial Examples”. In: (Feb. 2018). URL: https://openreview.net/forum?id=S18Su--CW.

[21] Chuan Guo et al. “Countering Adversarial Images using Input Transformations”. In: (Feb. 2018). URL: https://openreview.net/forum?id=SyJ7ClWCb.

[22] Claudio Michaelis et al. “Benchmarking Robustness in Object Detection: Autonomous Driving when Winter is Coming”. In: (2019), pp. 1–23. arXiv: 1907.07484. URL: http://arxiv.org/abs/1907.07484.

[23] Andrew Ilyas et al. “Adversarial Examples Are Not Bugs, They Are Features”. In: (May 2019). arXiv: 1905.02175 [stat.ML].

[24] D. Linsley et al. “What are the Visual Features Underlying Human Versus Machine Vision?” In: ICCV. 2017. ISBN: 9781538610343. DOI: 10.1109/ICCVW.2017.331. arXiv: 1701.02704.

[25] Robert Geirhos et al. “ImageNet-Trained CNNs Are Biased Towards Texture”. In: International Conference on Learning Representations c (2019), pp. 1–20. DOI: arXiv:1811. 12231v1. arXiv: 1811.12231.

[26] Daniel L. K. Yamins et al. “Performance-optimized hierarchical models predict neural responses in higher visual cortex”. In: Proceedings of the National Academy of Sciences 111.23 (2014), pp. 8619–8624. ISSN: 0027-8424. DOI: 10.1073/pnas.1403112111. arXiv: 0706.1062v1. URL: http://www.pnas.org/cgi/doi/10.1073/pnas.1403112111.

[27] Charles F Cadieu et al. “Deep neural networks rival the representation of primate IT cortex for core visual object recognition”. In: PLoS computational biology 10.12 (2014), e1003963.

[28] Seyed-Mahdi Khaligh-Razavi and Nikolaus Kriegeskorte. “Deep supervised, but not unsupervised, models may explain IT cortical representation”. en. In: PLoS Comput. Biol. 10.11 (Nov. 2014), e1003915.

[29] Umut Güçlü and Marcel A. J. van Gerven. “Deep Neural Networks Reveal a Gradient in the Complexity of Neural Representations across the Brain’s Ventral Visual Pathway”. In: The Journal of neuroscience 35.27 (2015), pp. 10005–10014. ISSN: 0270-6474. DOI: 10.1523/JNEUROSCI.5023-14.2015. arXiv: 1411.6422. URL: http://arxiv.org/abs/1411.6422%7B%5C%%7D0A http://dx.doi.org/10.1523/JNEUROSCI.5023-14.2015.

[30] Santiago A Cadena et al. “Deep convolutional models improve predictions of macaque V1 responses to natural images”. en. In: PLoS Comput. Biol. 15.4 (Apr. 2019), e1006897.

[31] Martin Schrimpf et al. “Brain-Score: Which Artificial Neural Network for Object Recognition is most Brain-Like?” en. Sept. 2018. URL: https://doi.org/10.1101/407007.

[32] Jonas Kubilius et al. “Brain-Like Object Recognition with High-Performing Shallow Recurrent ANNs”. In: Advances in Neural Information Processing Systems 32. Ed. by H. Wallach et al. Curran Associates, Inc., 2019, pp. 12805–12816. URL: http://papers.nips.cc/paper/9441-brain-like-object-recognition-with-high-performingshallow-recurrent-anns.pdf.

[33] Adam H Marblestone, Greg Wayne, and Konrad P Kording. “Toward an Integration of Deep Learning and Neuroscience”. en. In: Front. Comput. Neurosci. 10 (Sept. 2016), p. 94.

[34] William Lotter, Gabriel Kreiman, and David Cox. “Deep Predictive Coding Networks for Video Prediction and Unsupervised Learning”. In: (May 2016). arXiv: 1605.08104 [cs.LG].

[35] Aran Nayebi and Surya Ganguli. “Biologically inspired protection of deep networks from adversarial attacks”. In: (Mar. 2017). arXiv: 1703.09202 [stat.ML].

[36] Jordan Guerguiev, Timothy P Lillicrap, and Blake A Richards. “Towards deep learning with segregated dendrites”. en. In: Elife 6 (Dec. 2017).

[37] Demis Hassabis et al. “Neuroscience-Inspired Artificial Intelligence”. In: Neuron 95.2 (2017), pp. 245–258. ISSN: 10974199. DOI: 10.1016/j.neuron.2017.06.011. arXiv: 1404.7282.

[38] Grace W Lindsay and Kenneth D Miller. “How biological attention mechanisms improve task performance in a large-scale visual system model”. en. In: Elife 7 (Oct. 2018).

[39] Hanlin Tang et al. “Recurrent computations for visual pattern completion”. In: Proceedings of the National Academy of Sciences 115.35 (2018), pp. 8835–8840. ISSN: 0027-8424. DOI: 10.1073/pnas.1719397115.

[40] Zhe Li et al. “Learning From Brains How to Regularize Machines”. In: (Nov. 2019). arXiv: 1911.05072 [cs.LG].

[41] Hosein Hasani, Mahdieh Soleymani, and Hamid Aghajan. “Surround Modulation: A Bio-inspired Connectivity Structure for Convolutional Neural Networks”. In: NeurIPS NeurIPS (2019), pp. 15877–15888. URL: http://papers.nips.cc/paper/9719-surround-modulation-a-bio-inspired-connectivity-structure-for-convolutionalneural-networks.

[42] Fabian H Sinz et al. “Engineering a Less Artificial Intelligence”. en. In: Neuron 103.6 (Sept. 2019), pp. 967–979.

[43] Anthony M Zador. “A critique of pure learning and what artificial neural networks can learn from animal brains”. en. In: Nat. Commun. 10.1 (Aug. 2019), p. 3770.

[44] J Deng et al. “ImageNet: A large-scale hierarchical image database”. In: 2009 IEEE Conference on Computer Vision and Pattern Recognition. June 2009, pp. 248–255.

[45] D. H. Hubel and T. N. Wiesel. “Receptive Fields, Binocular Interaction and Functional Architecture in the Cat’s Visual Cortex”. In: Journal of Physiology (1962), pp. 106–154.

[46] J. P. Jones and L. A. Palmer. “An evaluation of the two-dimensional Gabor filter model of simple receptive fields in cat striate cortex”. In: Journal of Neurophysiology 58.6 (1987), pp. 1233–1258. ISSN: 0022-3077. DOI: 10.1152/jn.1987.58.6.1233. URL: http://www.physiology.org/doi/10.1152/jn.1987.58.6.1233.

[47] E.H. Adelson and J.R. Bergen. “Spatiotemporal energy models for the perception of motion”. In: Journal of the Optical Society of America A, Optics and image science 2.2 (1985), pp. 284–299. ISSN: 1084-7529. DOI: 10.1364/JOSAA.2.000284. URL:http://www.opticsinfobase.org/abstract.cfm?URI=josaa-2-2-284.

[48] David J. Heeger, E. P. Simoncelli, and J. Anthony Movshon. “Computational models of cortical visual processing”. In: Proceedings of the National Academy of Sciences of the United States of America 93.2 (1996), pp. 623–627. ISSN: 00278424. DOI: 10.1073/pnas.93.2.623.

[49] M. Carandini, D. J. Heeger, and J. A. Movshon. “Linearity and normalization in simple cells of the macaque primary visual cortex.” In: The Journal of neuroscience: the official journal of the Society for Neuroscience 17.21 (Nov. 1997), pp. 8621–44. ISSN: 0270-6474. URL: http://www.ncbi.nlm.nih.gov/pubmed/9334433.

[50] Maximilian Riesenhuber and Tomaso Poggio. “Hierarchical models of object recognition in cortex”. In: Nature Neuroscience 2.11 (1999), pp. 1019–1025. ISSN: 10976256. DOI: 10.1038/14819.

[51] Thomas Serre, Aude Oliva, and Tomaso Poggio. “A feedforward architecture accounts for rapid categorization”. en. In: Proc. Natl. Acad. Sci. U. S. A. 104.15 (Apr. 2007), pp. 6424–6429.

[52] Nicole C Rust et al. “Spatiotemporal Elements of Macaque V1 Receptive Fields”. In: Neuron 46 (2005), pp. 945–956. DOI: 10.1016/j.neuron.2005.05.021.

[53] Brett Vintch, J. A. Movshon, and E. P. Simoncelli. “A Convolutional Subunit Model for Neuronal Responses in Macaque V1”. In: Journal of Neuroscience 35.44 (2015), pp. 14829–14841. DOI: 10.1523/JNEUROSCI.2815-13.2015.

[54] Nicolas Papernot et al. “Distillation as a Defense to Adversarial Perturbations against Deep Neural Networks”. In: (Nov. 2015). arXiv: 1511.04508 [cs.CR].

[55] Mayank Singh et al. “Harnessing the Vulnerability of Latent Layers in Adversarially Trained Models”. In: (May 2019). arXiv: 1905.05186 [cs.LG].

[56] Cihang Xie et al. “Feature denoising for improving adversarial robustness”. In: Proceedings of the IEEE Conference on Computer Vision and Pattern Recognition. 2019, pp. 501–509.

[57] Eric Wong, Leslie Rice, and J Zico Kolter. “Fast is better than free: Revisiting adversarial training”. In: (Jan. 2020). URL: https://openreview.net/forum?id=BJx040EFvH.

[58] Dan Hendrycks et al. “AugMix: A Simple Data Processing Method to Improve Robustness and Uncertainty”. In: (Dec. 2019). arXiv: 1912.02781 [stat.ML].

[59] Evgenia Rusak et al. “Increasing the robustness of DNNs against image corruptions by playing the Game of Noise”. In: (Jan. 2020). arXiv: 2001.06057 [cs.CV].

[60] Ali Shafahi et al. “Adversarial training for free!” In: Advances in Neural Information Processing Systems 32. Ed. by H Wallach et al. Curran Associates, Inc., 2019, pp. 3358–3369.

[61] Eric Wong, Leslie Rice, and J Zico Kolter. “Fast is better than free: Revisiting adversarial training”. In: (Sept. 2020). URL: https://openreview.net/forum?id=BJx040EFvH.

[62] Dimitris Tsipras et al. “Robustness May Be at Odds with Accuracy”. Sept. 2019. URL: https://openreview.net/forum?id=SyxAb30cY7.

[63] Yash Sharma and Pin-Yu Chen. “Attacking the Madry Defense Model with L1-based Adversarial Examples”. In: (Oct. 2017). arXiv: 1710.10733 [stat.ML].

[64] Lukas Schott et al. “Towards the first adversarially robust neural network model on MNIST”. In: (May 2018). arXiv: 1805.09190 [cs.CV].

[65] Cihang Xie et al. “Mitigating Adversarial Effects Through Randomization”. Feb. 2018. URL: https://openreview.net/forum?id=Sk9yuql0Z.

[66] Adnan Siraj Rakin, Zhezhi He, and Deliang Fan. “Parametric Noise Injection: Trainable Randomness to Improve Deep Neural Network Robustness against Adversarial Attack”. In: (Nov. 2018). arXiv: 1811.09310 [cs.LG].

[67] Joonho Lee et al. “ProbAct: A Probabilistic Activation Function for Deep Neural Networks”. In: (May 2019). arXiv: 1905.10761 [cs.LG].

[68] Anish Athalye, Nicholas Carlini, and David Wagner. “Obfuscated Gradients Give a False Sense of Security: Circumventing Defenses to Adversarial Examples”. In: (Feb. 2018). arXiv: 1802.00420 [cs.LG].

[69] Alex Krizhevsky. “Learning Multiple Layers of Features from Tiny Images”. In: (Apr. 2009). URL: http://citeseerx.ist.psu.edu/viewdoc/download?doi=10.1.1.222.9220&rep&#x003D;rep1&type=pdf.

[70] Adam Paszke et al. “Automatic differentiation in PyTorch”. In: (Oct. 2017). URL: https://openreview.net/forum?id=BJJsrmfCZ.

[71] Saining Xie et al. “Aggregated Residual Transformations for Deep Neural Networks”. In: (Nov. 2016). arXiv: 1611.05431 [cs.CV].

[72] Gao Huang et al. “Densely Connected Convolutional Networks”. In: (Aug. 2016). arXiv: 1608.06993 [cs.CV].

[73] Forrest N Iandola et al. “SqueezeNet: AlexNet-level accuracy with 50x fewer parameters and <0.5mb model size”. In: (Feb. 2016). arXiv: 1602.07360 [cs.CV].

[74] Ningning Ma et al. “ShuffleNet V2: Practical Guidelines for Efficient CNN Architecture Design”. In: (July 2018). arXiv: 1807.11164 [cs.CV].

[75] Mingxing Tan et al. “MnasNet: Platform-Aware Neural Architecture Search for Mobile”. In: (July 2018). arXiv: 1807.11626 [cs.CV].

[76] Logan Engstrom et al. Robustness (Python Library). 2019. URL: https://github.com/MadryLab/robustness.

[77] Jeremy Freeman et al. “A functional and perceptual signature of the second visual area in primates”. en. In: Nat. Neurosci. 16.7 (July 2013), pp. 974–981.

[78] Russell L. De Valois, E. W. Yund, and Norva Hepler. “The orientation and direction selectivity of cells in macaque visual cortex”. In: Vision Research 22 (1982), pp. 531–544.

[79] Russell L. De Valois, Duane G. Albrecht, and Lisa G. Thorell. “Spatial Frequency Selectivity of Cells in Macaque Visual Cortex”. In: Vision Research 22 (1982), pp. 545–559.

[80] Dario L Ringach. “Spatial Structure and Symmetry of Simple-Cell Receptive Fields in Macaque Primary Visual Cortex”. In: Journal of Neurophysiology 88 (2002), pp. 455–463.

[81] Yasmine El-Shamayleh et al. “Visual response properties of V1 neurons projecting to V2 in macaque”. In: Journal of Neuroscience 33.42 (2013), pp. 16594–16605. ISSN: 02706474. DOI: 10.1523/JNEUROSCI.2753-13.2013.

[82] W. R. Softky and C. Koch. “The highly irregular firing of cortical cells is inconsistent with temporal integration of random EPSPs”. In: Journal of Neuroscience 13.1 (1993), pp. 334–350. ISSN: 02706474. DOI: 10.1523/jneurosci.13-01-00334.1993.

[83] Diederik P Kingma and Max Welling. “Auto-Encoding Variational Bayes”. In: (Dec. 2013). arXiv: 1312.6114v10 [stat.ML].

[84] Nicholas Carlini et al. “On Evaluating Adversarial Robustness”. In: (Feb. 2019). arXiv: 1902.06705v2 [cs.LG].

[85] Nikolaus Kriegeskorte, Marieke Mur, and Peter Bandettini. “Representational similarity analysis - connecting the branches of systems neuroscience”. en. In: Front. Syst. Neurosci. 2 (Nov. 2008), p. 4.

[86] R. Rajalingham, K. Schmidt, and James J. DiCarlo. “Comparison of Object Recognition Behavior in Human and Monkey”. In: Journal of Neuroscience 35.35 (2015), pp. 12127–12136. ISSN: 0270-6474. DOI: 10.1523/JNEUROSCI.0573-15.2015. URL: http://www.jneurosci.org/cgi/doi/10.1523/JNEUROSCI.0573-15.2015.

[87] Cristopher M Niell and Michael P Stryker. “Highly selective receptive fields in mouse visual cortex.” In: The Journal of neuroscience 28.30 (July 2008), pp. 7520–36. ISSN: 1529-2401. DOI: 10.1523/JNEUROSCI.0623-08.2008. URL: http://www.ncbi.nlm.nih.gov/pubmed/18650330.

[88] Pieter R Roelfsema. “Cortical algorithms for perceptual grouping.” In: Annual review of neuroscience 29.March (Jan. 2006), pp. 203–27. ISSN: 0147-006X. DOI: 10.1146/annurev.neuro.29.051605.112939. URL: http://www.ncbi.nlm.nih.gov/pubmed/16776584.

[89] Alexander Berardino et al. “Eigen-Distortions of Hierarchical Representations”. In: (Oct. 2017). arXiv: 1710.02266 [cs.CV].

[90] Jack Lindsey et al. “A Unified Theory of Early Visual Representations from Retina to Cortex through Anatomically Constrained Deep CNNs”. In: ICLR (2019), pp. 1–17. DOI: 10.1101/511535. arXiv: 1901.00945. URL: http://arxiv.org/abs/1901.00945.

[91] Daniel L K Yamins and James J DiCarlo. “Using goal-driven deep learning models to understand sensory cortex”. en. In: Nat. Neurosci. 19.3 (Mar. 2016), pp. 356–365.

[92] Alexander Mathis et al. “DeepLabCut: markerless pose estimation of user-defined body parts with deep learning”. In: Nature Neuroscience (2018). ISSN: 1097-6256. DOI: 10.1038/s41593-018-209-y. arXiv: 1804.03142. URL: http://www.nature.com/articles/s41593-018-0209-y.

[93] Yann LeCun, Yoshua Bengio, and Geoffrey E. Hinton. “Deep learning”. In: Nature 521.7553 (2015), pp. 436–444. ISSN: 14764687. DOI: 10.1038/nature14539. arXiv: 1807.07987.

[94] T. Marques, M. Schrimpf, and J. J. DiCarlo. “Hierarchical neural network models that more closely match primary visual cortex tend to better explain higher level visual cortical responses”. In: Cosyne. 2020.

[95] Alexey Kurakin, Ian Goodfellow, and Samy Bengio. “Adversarial examples in the physical world”. In: (July 2016). arXiv: 1607.02533 [cs.CV].

[96] Maria-Irina Nicolae et al. “Adversarial Robustness Toolbox v1.2.0”. In: CoRR 1807.01069 (2018). URL: https://arxiv.org/pdf/1807.01069.

[97] Florian Tramer et al. “On Adaptive Attacks to Adversarial Example Defenses”. In: (Feb. 2020). arXiv: 2002.08347 [cs.LG].

[98] Jonas Rauber et al. “Foolbox Native: Fast adversarial attacks to benchmark the robustness of machine learning models in PyTorch, TensorFlow, and JAX”. In: Journal of Open Source Software 5.53 (2020), p. 2607. DOI: 10.21105/joss.02607. URL: https://doi.org/10.21105/joss.02607.

[99] Matthew D. Zeiler and Rob Fergus. “Visualizing and Understanding Convolutional Networks”. In: arXiv (Nov. 2013). ISSN: 16113349. DOI: 10.1007/978-3-319-10590-153. arXiv: 1311.2901. URL: http://link.springer.com/10.1007/978-3-319-10590-1%7B%5C_%7D53%7B%5C%%7D5Cn http://arxiv.org/abs/1311.2901%7B%5C%%7D5Cnpapers3://publication/uuid/44feb4b1-873a-4443-8baa-1730ecd16291%20 http://arxiv.org/abs/1311.2901.

[100] Chris Olah et al. “An Overview of Early Vision in InceptionV1”. In: Distill (2020). https://distill.pub/2020/circuits/early-vision. DOI: 10.23915/distill.00024.002.

[101] Ian Nauhaus et al. “Orthogonal micro-organization of orientation and spatial frequency in primate primary visual cortex”. In: Nature Neuroscience 15.12 (2012), pp. 1683–1690. ISSN: 10976256. DOI: 10.1038/nn.3255.

